# Eomesodermin is functionally conserved between zebrafish and mouse in spite of different mutant phenotypic severities, and controls left/right organiser formation via interlocking feedforward loops

**DOI:** 10.1101/2020.10.02.324244

**Authors:** Conor D. Talbot, Mark D. Walsh, Stephen J. Cutty, Randa Elsayed, Ashley E. E. Bruce, Fiona C. Wardle, Andrew C. Nelson

**Affiliations:** School of Life Sciences, Gibbet Hill Campus, University of Warwick, Coventry, CV4 7AL, UK; Randall Centre for Cell and Molecular Biophysics, New Hunt’s House, Guy’s Campus, King’s College London, SE1 1UL, UK; Warwick Medical School, Gibbet Hill Campus, University of Warwick, Coventry, CV4 7AL, UK; Department of Cell and Systems Biology, University of Toronto, Toronto, ON M5S 3G5, Canada

## Abstract

The T-box family transcription factor Eomesodermin (Eomes) is present in all vertebrates, with many key roles in the developing mammalian embryo and immune system. Homozygous Eomes mutant mouse embryos exhibit early lethality due to defects in both the embryonic mesendoderm and the extraembryonic trophoblast cell lineage. In contrast, zebrafish lacking the predominant Eomes homologue A (Eomesa) do not suffer complete lethality and can be maintained. This suggests fundamental differences in either the molecular function of Eomes orthologues or the molecular configuration of processes in which they participate. To explore these hypotheses we initially analysed the expression of distinct Eomes isoforms in various cell mouse types. Next we compared the functional capabilities of these murine isoforms compared to zebrafish Eomesa. These experiments provided no evidence for functional divergence. Next we examined the functions of zebrafish Eomesa and other T-box family members expressed in early development, as well as its paralogue Eomesb. Though Eomes is a member of the Tbr1 subfamily we found strong evidence for functional redundancy without complete functional equivalence with the Tbx6 subfamily member Tbx16, known to be absent from eutherians and other mammals. Finally, we analysed the ability of Eomesa to induce zebrafish left-right organiser progenitors (known as dorsal forerunner cells) known to be positively regulated by *vgll4l*, a gene we had previously shown to be repressed by Eomesa. Here we demonstrate that Eomesa indirectly upregulates *vgll4l* expression via interlocking feedforward loops, suggesting a role in establishment of left/right asymmetry. Overall these findings demonstrate conservation of Eomes molecular function and participation in similar processes, but differential requirements across evolution due to the expanded complement of T-box factors in teleosts. Our analyses also provide insights into the role of Eomesa in left-right organiser formation in zebrafish.

**Author summary:** Recent studies provide evidence for both shared and unique molecular pathways controlling early development in different vertebrate organisms. The transcription factor Eomesodermin plays essential roles during mammalian development and has potent functional capabilities in zebrafish embryos yet is seemingly dispensable for viability. Here we compared functional contributions of the predominant zebrafish Eomesodermin homologue - Eomesa - with multiple isoforms of mouse Eomesodermin. We found them to be functionally indistinguishable in the early embryo. Surprisingly, a distant relative of Eomesodermin in the zebrafish embryo – Tbx16 – which is absent in mammals shows a degree of functional overlap with Eomesodermin, underscoring the different evolutionary requirements for Eomesodermin in teleosts and mammals. Finally, we demonstrate that Eomesodermin differentially regulates the transcriptional cofactor *vgll4l*, which plays key roles in formation of the zebrafish left/right organiser, at different stages of development. Thus, our work demonstrates the molecular function of Eomesodermin is likely to be conserved throughout vertebrate evolution despite differences in mutant phenotypes, and reveals regulatory mechanisms controlling left/right asymmetric patterning.

## Introduction

T-box transcription factors (TFs) are an ancient family of transcriptional regulators with diverse roles in development and disease [1]. Eomesodermin (Eomes) belongs to the Tbr1 subfamily of T-box TFs, consisting of similarly sized N- and C-terminal domains (NTD and CTD) flanking a central DNA binding domain known as the T-box. Amongst the species where *Eomes* is best studied are mouse and zebrafish [2]. Mice have a single copy of *Eomes*, whereas zebrafish owing to the whole genome duplication in the teleost lineage have two paralogous genes, *eomesa* and *eomesb* [3]. During mouse embryogenesis *Eomes* plays essential roles in trophectoderm [4, 5], in the primitive streak for epithelial-to-mesenchymal transition, mesoderm migration and specification of definitive endoderm and cardiac mesoderm during gastrulation [6, 7]. Additionally *Eomes* acts in the visceral endoderm to control anterior-posterior axis identity [8] and later has key functions in cortical neuron progenitors [9]. It is also expressed in progenitors of the left/right organiser known in mammals as the node, and is required for correct formation of the node suggesting a potential role in establishing left/right asymmetry [6, 7]. In zebrafish Eomesa also plays a role in mesenoderm formation, being able to induce ectopic endoderm if overexpressed with essential interacting factors [10], and is also sufficient to induce dorsal mesoderm markers and represses ectoderm gene expression in early development [11, 12]. Eomesa is also sufficient to induce progenitors of the left/right organiser, known as dorsal forerunner cells (DFCs) in zebrafish [10]. However, we recently found that Eomesa represses expression of the transcriptional cofactor *vgll4l* [12], a key positive regulator of DFC proliferation, survival and function [13]. Here we further investigate these paradoxical findings.

Mouse *Eomes* and zebrafish *eomesa* display similar expression domains during early development [4, 14-18]. However, surprisingly endoderm, cardiac mesoderm and axial patterning proceed normally in *eomesa* loss-of-function mutants [17]. This observation cannot be explained simply by rescue by *eomesb*, which is not co-expressed with *eomesa* in early development [12, 19]. The extent to which Eomes functional activities are conserved between zebrafish and mouse remains unknown.

One possibility is that these distinct loss-of-function phenotypes could potentially be due to functional diversification during evolution. The process of alternative splicing (AS) allows a single gene to give rise to multiple isoforms with different functional characteristics. The prevalence of AS has expanded across evolutionary time, allowing increased proteome diversity out of proportion with gene number [20]. For example, only ∼25% of nematode genes have alternative isoforms compared to >90% in human [21, 22]. AS leading to functional diversification may account for altered functions of Eomes between species. However, it is also possible that differential requirement for Eomes is due to functional redundancy owing to altered complements of T-box factors in different vertebrate evolutionary lineages. The most ancient T-box factor *Brachyury* is present in several non-metazoan lineages, however, the T-box family is considerably expanded in Metazoa, reflecting its developmental importance [23]. Additionally, the complement of T-box factors has varied across vertebrate evolution, with gain or loss of individual factors in certain lineages. For example, the Tbx6 subfamily member *tbx16* is present in fish, frogs and birds but is lost in mammals. The T-box domain itself directly binds DNA in a sequence-specific manner. Genome-wide profiling of multiple T-box factors including Eomes, Tbx16, Tbx6 and Brachyury in zebrafish, mice, *Xenopus* and human has revealed they bind most frequently to variants of an eight to nine base pair core consensus of (T)TVRCACHT, interchangeably allowing occupancy of different T-box factors at the same genomic sites e.g. [12, 24-33]. T-box factors therefore often exhibit redundancy through regulation of the same target genes through the same *cis*-regulatory modules.

We therefore sought to answer three key questions: 1. Are zebrafish and mouse *Eomes* genes functionally equivalent? 2. What is the basis for the observed differences in severity of loss-of-function phenotypes between mouse and zebrafish? and; 3. How can Eomesa promote DFC formation while repressing the key DFC regulator *vgll4l*?

Our analyses suggest that the molecular function of *Eomes* is highly conserved throughout vertebrate evolution. Our data also reveal that while alternative splicing of mouse *Eomes* transcript occurs at exon 6, functionally the encoded proteins were indistinguishable. We found that Eomesa and Tbx16 share overlapping functions and capabilities in the presumptive endoderm, suggesting that phenotypic rescue by Tbx16 may explain *eomesa* mutant viability. Finally, we found that Eomesa acts within interlocking feedforward loops to both repress *vgll4l* and activate it indirectly via the essential SOX family transcription factor Sox32. Our results therefore advance our understanding of T-box factor functional conservation during early vertebrate embryogenesis, and the gene regulatory networks controlling left/right organiser formation.

## Results

### Eomes isoforms, conservation and expression

In mouse the three annotated *Eomes* transcripts give rise to three structurally distinct isoforms including the full length (FL) product, a splice variant having an alternative splice acceptor site within exon 6 leading to loss of a 19 amino acid variant region (ΔVR), and a transcript with an alternative mRNA 3’ cleavage site leading to loss of exon 6 and its encoded CTD (ΔCTD) (Fig 1A). The highly conserved VR sequence is known to be phosphorylated at three amino acid residues in mouse spleen and kidney (Fig 1A) [34]. The internal exon 6 splicing event emerged in the tetrapod lineage through a synonymous single nucleotide change in an arginine codon (CG>AG) introducing a splice acceptor sequence.

**Figure 1.**
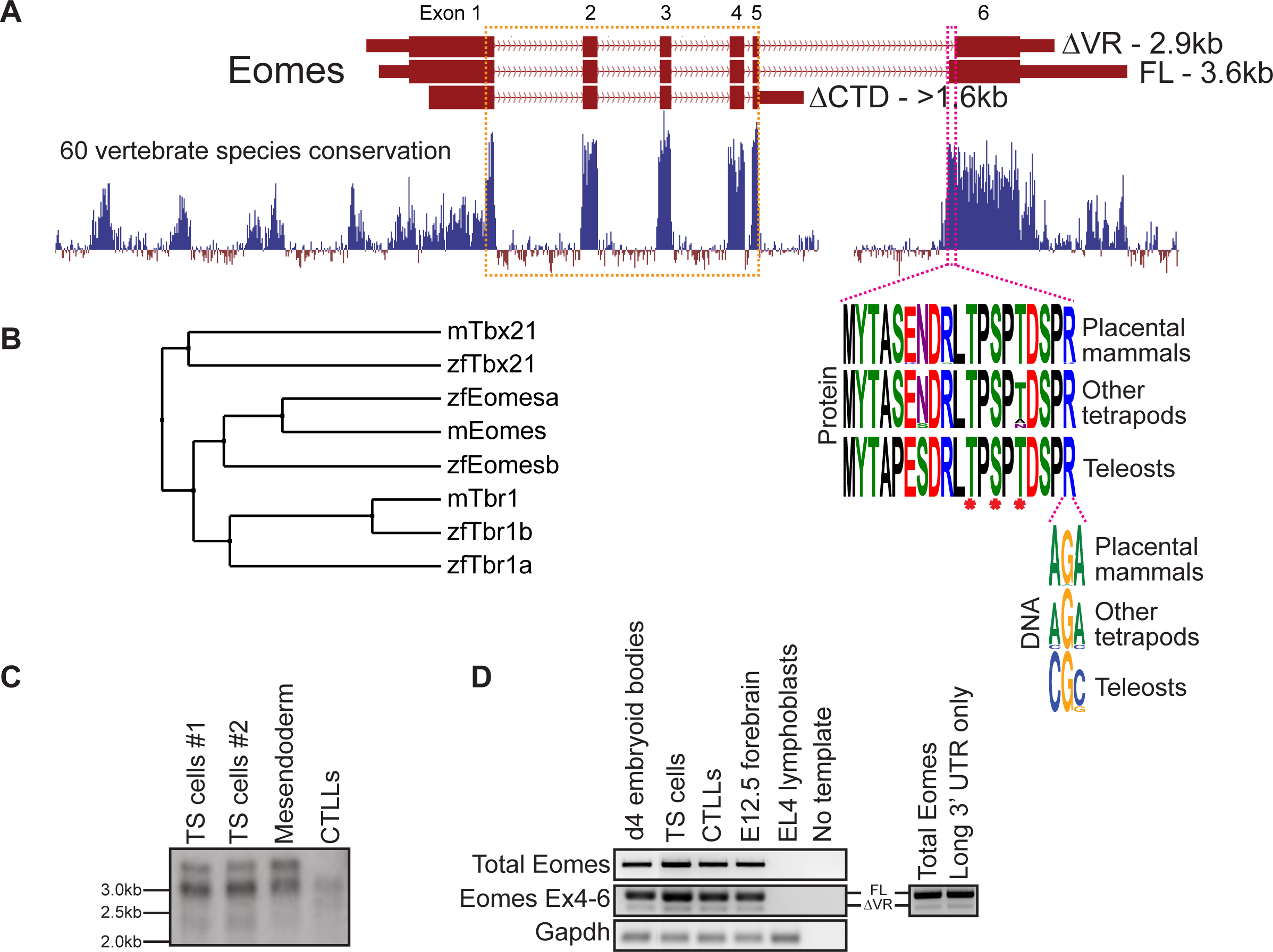
Mouse *Eomes* has multiple isoforms, including a mammalian-specific alternative splicing event. (A) Gene model with conservation track and sequence logos for variant region. Annotated transcript sizes are indicated, as well as amino acid conservation of the VR between placental mammals, other tetrapods and teleosts, and the variation within the terminal VR arginine codon. The T-box is outlined in orange and the VR in pink. Asterisks indicates known phosphorylated amino acid residues. (B) BLOSUM62 average distance evolutionary tree of the Tbr1 subfamily showing relationships between mouse and zebrafish genes. (C) Northern blot showing Eomes transcripts in different cell types using a probe against the T-box. Data for two independent trophoblast stem (TS) cell lines are shown. Mesendoderm is P19Cl6 cells after 4 days of DMSO induced differentiation. CTLLs are IL-2-dependent T-cell lymphocytes derived from ATCC TIB-214. (D) RT-PCR showing relative levels of FL and ΔVR isoforms (left), and nested PCR showing FL/ΔVR ratio for long 3’ UTR transcripts (right). Day 4 differentiated embryoid bodies contain cells mimicking embryonic endoderm. CTLLs are IL-2-dependent T-cell lymphocytes derived from ATCC TIB-214. EL4 cells are a negative control for *Eomes* expression. *Gapdh* is a loading control.

Because the ΔCTD transcript annotation has an incomplete 5’ end, it remains unclear whether it encodes the entire NTD. However, our RT-PCR analysis using primers located in the ΔCTD 3’ UTR and at the FL/ΔVR start codon suggests that exon 1 coding information is intact (not shown). The CTD encoded by exon 6 has been shown to function in transcriptional activation [35], suggesting that the ΔCTD isoform is likely to be functionally compromised in comparison to FL and ΔVR isoforms. Consistent with this, the CTD is more highly conserved than the NTD (Fig 1A). Functional differences between FL and ΔVR isoforms, however, are yet to be examined. Both *eomesa* transcripts identified in zebrafish encode the same protein [11]. Relatively little is known about the single annotated zebrafish *eomesb* transcript, which appears to be more divergent from the ancestral gene (Fig 1B).

Murine *Eomes* is expressed in numerous cell types including trophoblast stem cells (TSCs), mesendoderm, and T lymphocytes. To identify *Eomes* transcripts we initially performed Northern blot analysis (Fig 1C). Transcript sizes corresponding to all three annotated isoforms were detectable but the ΔCTD transcript was underrepresented. The FL and ΔVR annotations display different 3’ UTR lengths. To test whether the two distinct upper bands detected by Northern blot correspond to alternative exon 6 splicing events or merely different UTR lengths we next performed nested PCR (Fig 1D). We found that the long 3’ UTR is associated with both the FL and ΔVR coding isoforms. Strikingly, the ratio of FL/ΔVR is the same for both total Eomes and the long 3’ UTR transcripts. The abundance of the different coding transcripts therefore appears to be independent of UTR length. Further analysis through cloning *Eomes* intron5/exon6 to replace intron2/exon3 of the human *HBB* gene in an expression construct followed by transfection into HeLa cells revealed that the ratio of FL/ΔVR splicing is consistent with the wild type *Eomes* gene, suggesting that the low levels of the ΔVR isoform are due to weaker splicing consensus sequences, thus favouring the FL isoform (Fig S1). However, analysis of Eomes proteins by Western blot indicates that various N-terminal truncations occur which cannot be accounted for by the annotated coding transcripts (Fig S2). We conclude that FL is clearly the most abundant of the three annotated coding isoforms. Importantly, this predominant isoform expressed by mouse cells corresponds to the single *eomesa* isoform in zebrafish.

#### Both mouse FL and ΔVR isoforms are functionally equivalent to zebrafish eomesa in early development

Zebrafish *eomesa* loss of function causes less severe phenotypes in compared with the dramatic defects observed in *Eomes* mutant embryos. To further explore mouse and zebrafish Eomes functional capabilities we overexpressed either mouse *Eomes* FL or ΔVR mRNAs in zebrafish embryos for comparison with those overexpressing zebrafish *eomesa*.

Zebrafish Eomesa represses ectoderm genes such as *vgll4l* and *zic3* during blastula stages and activates mesendoderm genes including organizer markers *noto* and *chd* at the onset of gastrulation [11, 12]. Injecting equimolar quantities of zebrafish *eomesa* mRNA, or mouse FL or ΔVR Eomes isoforms at the one-cell stage, we found that each was equally able to repress *vgll4l* and *zic3* at 4 hours post-fertilisation (h.p.f.) and induce *noto* and *chd* at 6 h.p.f. (Fig 2A-D). We conclude that both FL and ΔVR mouse Eomes isoforms are functionally equivalent to zebrafish Eomesa.

**Figure 2.**
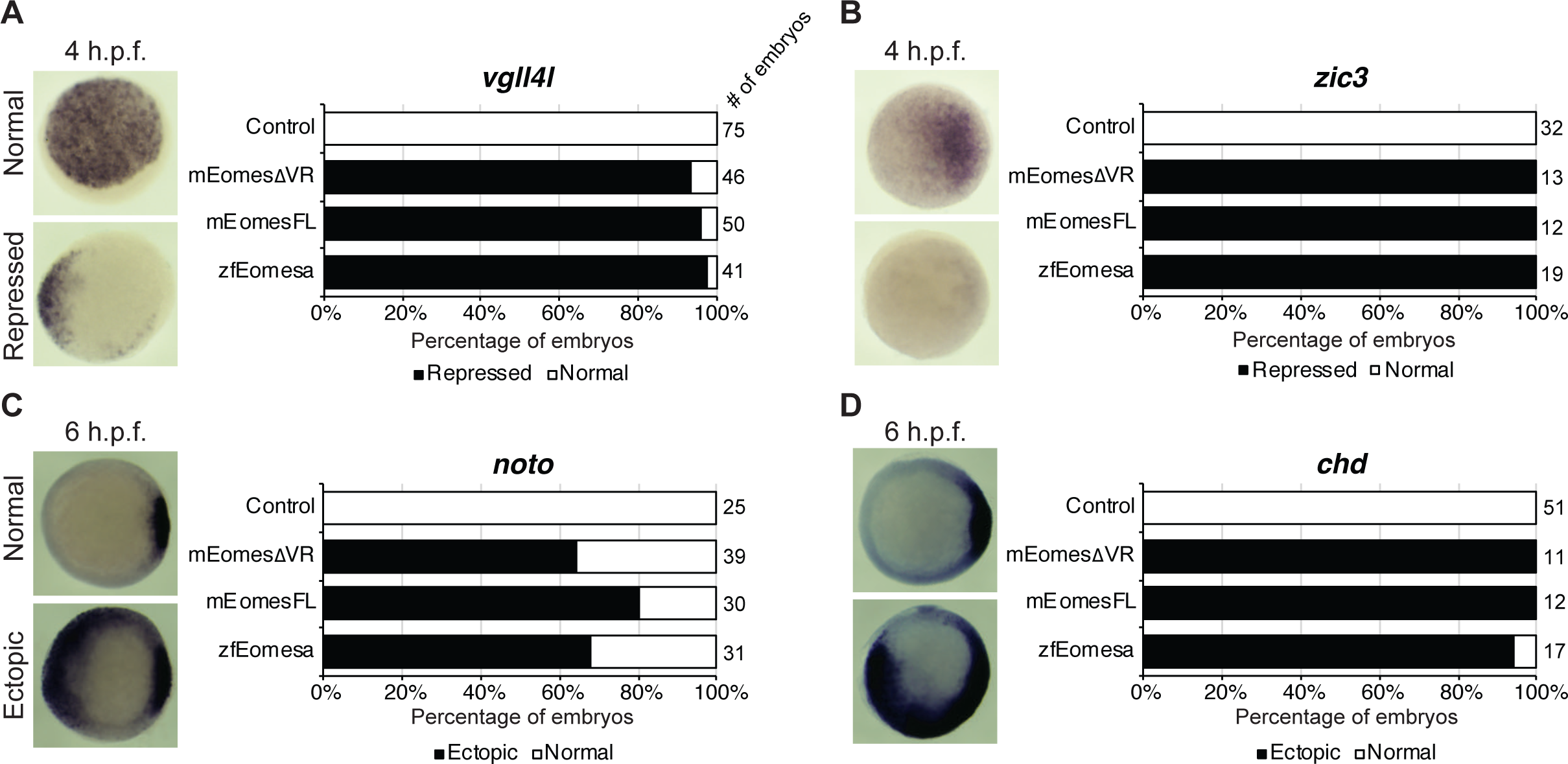
Both FL and ΔVR isoforms of mouse Eomes are functionally equivalent to zebrafish Eomesa in the early embryo. WISH analysis of ectoderm markers *vgll4l* and *zic3* in sphere stage (4 h.p.f.) embryos (A-B) or organiser markers *noto* and *chd* (C-D) at shield stage (6 h.p.f.). Embryos have been injected at the one cell stage to overexpress either mouse EomesFL, EomesΔVR or zebrafish Eomesa. N = 2. Total numbers of embryos scored per condition are indicated. Representative images of expression patterns per gene per category are shown. Animal views; dorsal to the right.

### Eomesa and non-mammalian T-box factor Tbx16 redundantly regulate mixl1 expression at the initiation of zebrafish endoderm formation

Since results above strongly suggest mouse Eomes is functionally equivalent to zebrafish Eomesa, do *eomesa* mutants have comparatively mild defects due to functional redundancy with other T-box factors? *Eomesb* is not appreciably expressed during early zebrafish development (Fig 3A), and is not upregulated in *eomesa* mutants [12] thus it seems unlikely that it compensates for loss of Eomesa. We recently identified a key role for the non-mammalian T-box factor, Tbx16 in endoderm formation, with Tbxta having a more minor role [33]. Both of these factors show zygotic upregulation concomitant with declining *eomesa* mRNA levels and prior to expression of key markers of presumptive endoderm (e.g. *gata5* and *mixl1*), and endoderm (e.g. *gata5, sox32* and *sox17*; Fig 3A), thus may compensate for the early loss of expression of such markers in MZ*eomesa* mutants [17].

**Figure 3.**
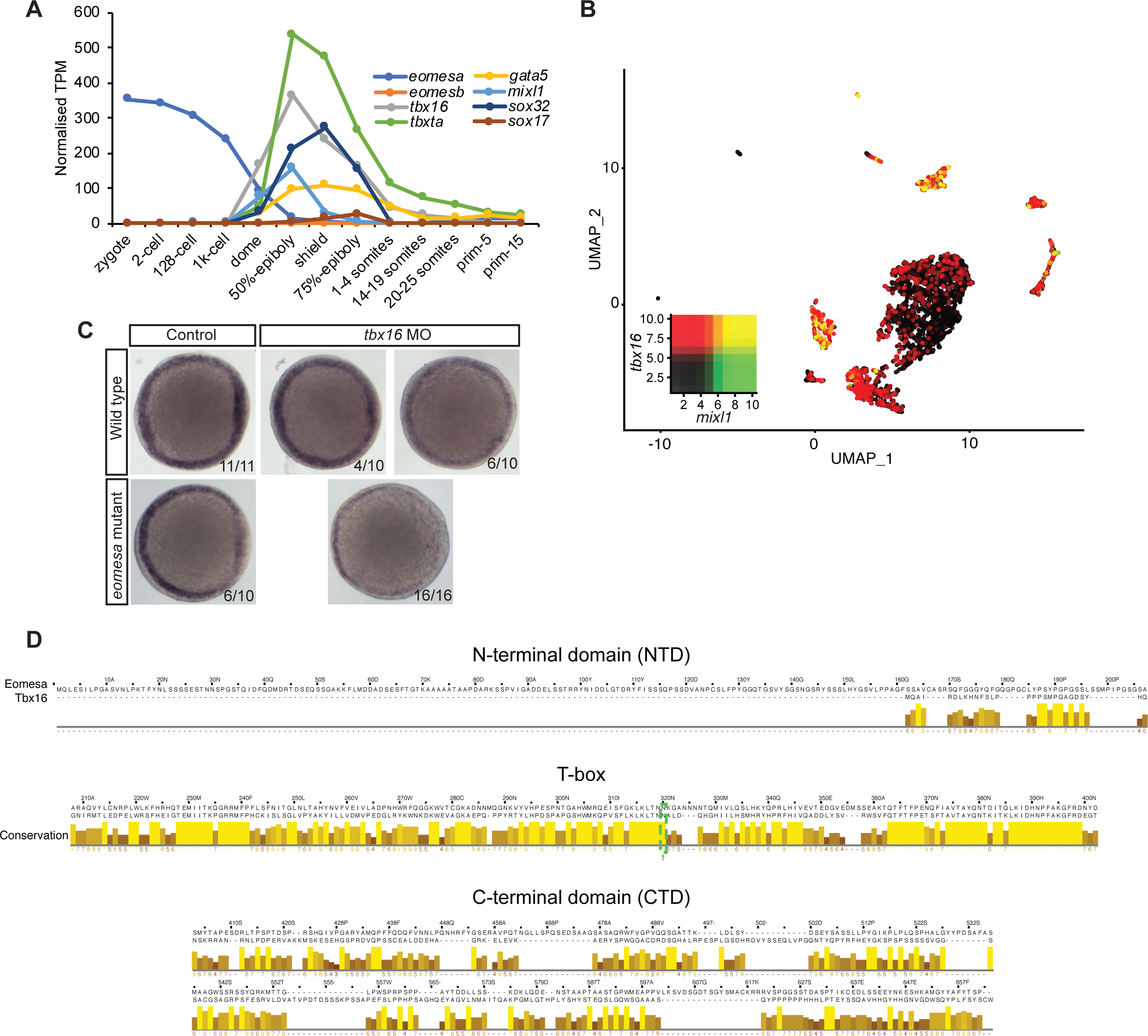
Eomesa and Tbx16 are redundantly required for *mixl1* expression in the presumptive endoderm. (A) Timing of expression of T-box factors (*eomesa, eomesb, tbxta, tbx16*) under study and presumptive endoderm (*mixl1, gata5, sox32*) and endoderm markers (*gata5, sox32, sox17*) indicated by bulk RNA-seq data from [81]. Gene expression is shown as transcripts per million (TPM). Stages are as defined by [82]. (B) UMAP clustering analysis of single-cell RNA-seq data for whole shield stage (6 h.p.f.) zebrafish embryos indicating co-expression of *tbx16* and *mixl1* [36]. Heatmap insets indicate overall expression levels per gene and co-expression. Overlapping expression is shown in yellow. (C) WISH analysis of *mixl1* at germ ring – shield stage (5.7-6 h.p.f.) zebrafish embryos in wild type or *eomesa* mutant embryos (see Methods for information on genotype) with and without Tbx16 morpholino knockdown. Total numbers of embryos and fractions as categorised are indicated. Animal views; dorsal to the right. (D) Jalview alignment of zebrafish FL Eomesa and Tbx16 proteins separated by domain. The conserved N amino acid at Eomesa position 320 is outlined in green. Conservation scores indicate similarity of physico-chemical properties of aligned amino acids

Moreover, single-cell RNA-seq experiments demonstrate that *tbx16* is robustly co-expressed with the critical endodermal regulator *mixl1* at 6 h.p.f., strongly suggesting that Tbx16 upregulates *mixl1* expression to initiate endoderm specification [33, 36] (Fig 3B). To test whether Tbx16 functions redundantly with Eomesa during endoderm formation next we performed antisense morpholino knockdown of Tbx16 in wild type and *eomesa* mutant embryos. We found that while *mixl1* expression is reduced on loss of Eomesa or Tbx16 alone, loss of both TFs leads to more striking loss of *mixl1* (Fig 3C). Eomesa and Tbx16 therefore collaboratively activate *mixl1* expression, strongly suggesting that Tbx16 relieves the requirement for Eomesa in zebrafish endoderm formation.

### Functional redundancy of Eomesa with co-expressed and related T-box factors is not driven by molecular equivalence

Tbx16 and Eomesa lack significant sequence homology, especially outside the T-box domain (Fig 3D). However, *Xenopus* Eomes and its Tbx16 orthologue VegT have been suggested to display similar specificity in part due to a single shared asparagine residue within the T-box, rendering them functionally distinct from the Tbxta orthologue Xbra, which has a lysine in the equivalent position [37] (Fig 3D). We therefore sought to address the following questions: 1. Do Tbx16, Tbxta and/or Eomesb have functional equivalence to Eomesa in early development; 2. Is the T-box asparagine residue critical for Eomesa function; 3. Are key Eomesa functions dependent on the highly conserved CTD or relatively poorly conserved NTD?

We injected equimolar quantities of mRNA corresponding to each wild type or mutant gene and assessed the effect on dorsal mesoderm marker *noto* and DFC/endoderm marker *sox32*. Deletion of NTD or CTD ablated Eomesa ability to induce *noto* expression, while N320K mutation had no significant effect (Fig 4A). All other T-box factors failed to produce a consistent or compelling induction of ectopic *noto* (Fig 4A). Consistent with previous results, Eomesa overexpression led to ectopic *sox32* expression in the outer marginal cells indicative of increased numbers of DFCs but not endoderm [10] (Fig 4B). NTD and especially CTD deletions markedly reduced ectopic *sox32* induction, however, N320K mutation had little effect (Fig 4B). Overexpression of *tbx16, tbxta and eomesb* did not lead to ectopic *sox32* induction highlighting functional distinctions with *eomesa* (Fig 4B).

**Figure 4.**
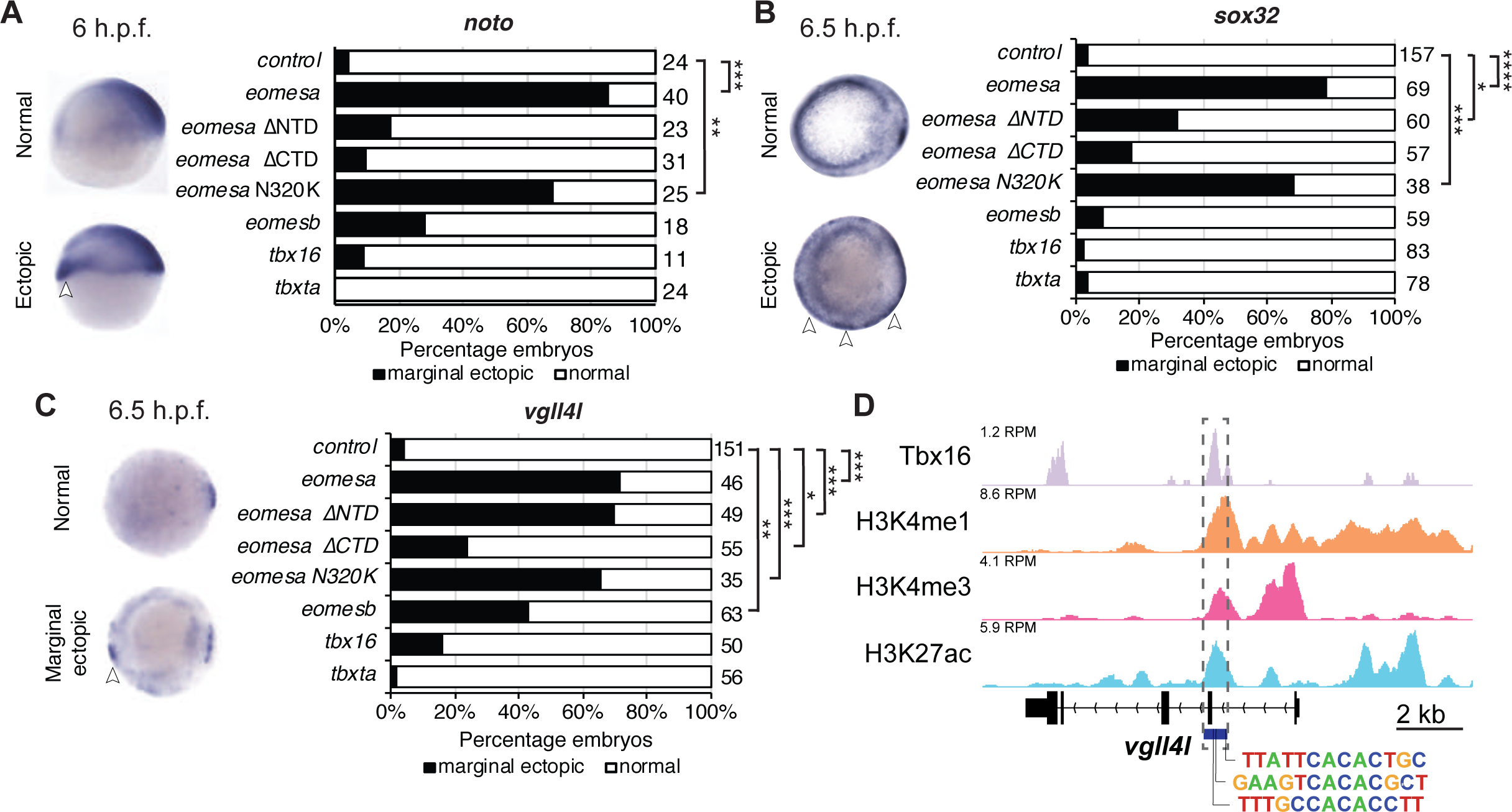
Eomesa is a more potent inducer of endoderm, organiser and dorsal forerunner cell markers than other T-box factors. (A-C) WISH analysis of dorsal mesoderm marker *noto* (A) and dorsal forerunner cell markers *sox32* (B) and *vgll4l* (C) on overexpression of various wild type and mutant T-box factors. mRNAs injected at the one cell stage; WISH performed at stages as indicated. N = 2. Total numbers of embryos scored per condition are indicated. Representative images of expression patterns per gene per category are shown. Panel A lateral views; panel B and C animal views; dorsal to the right. Arrowheads indicate ectopic expression. Fisher’s Exact two-tailed probability test P values: * *P* ≤ 5×10^−4^; ** *P* ≤ 5×10^−6^; *** *P* ≤ 5×10^−12^; **** *P* ≤ 5×10^−30^. (D) ChIP-seq data at 75-85% epiboly (8-8.5 h.p.f.) indicating Tbx16 binding within the *vgll4l* promoter [33, 83]. Scale is reads per million reads (RPM). Putative T-box binding sites identified using JASPAR are indicated [84].

Though Eomesa represses expression of *vgll4l* during blastula stages, during gastrulation *vgll4l* is expressed in DFCs, and has recently been identified as a key regulator of DFC proliferation, survival and function [13]. Repression of *vgll4l* at later stages in DFCs would consequently be inconsistent with promoting DFC formation. We therefore also tested whether the T-box factors of interest affect *vgll4l* expression consistent with roles in DFC formation. *Vgll4l* expression showed similar induction to *sox32* at the margin by both wild type Eomesa and ΔNTD and N320K Eomesa (Fig 4C). We have previously shown by ChIP-seq that at sphere stage Eomesa binds in the first intron of *vgll4l* [12]. Analysis of Tbx16 ChIP-seq data at 80-85% epiboly [33] also shows Tbx16 binding at close matches to the known T-box consensus sequence within *vgll4l* intron 1, suggesting Tbx16 does have the potential to directly participate in regulation of *vgll4l* during gastrulation (Fig 4D). However, *tbx16* overexpression suggests that it is not individually sufficient to strongly drive ectopic *vgll4l* expression (Fig 4C).

Overall this suggests that these T-box TFs do not have equivalent function in early development, that the previously reported N/K amino acid difference between Eomesa/Tbx16 and Tbxta does not appreciably influence specificity and function in this context, and that deletion of the relatively poorly conserved Eomesa NTD does not result in complete loss of function.

### Tbx16 and eomesa overexpression do not equivalently induce endoderm fate in concert with mixl1 and gata5

Given Eomesa is established to induce ectopic endoderm/DFC marker *sox32* as part of a complex with Mixl1 and Gata5, we extended our study to test whether Tbx16 can similarly induce ectopic endoderm amongst cells at the animal pole through co-overexpression with *mixl1* and *gata5*. We note that while *tbx16* is substantially co-expressed with *sox32* at the onset of gastrulation, *tbxta* is not (Fig 5A-B). This suggests that Tbx16 but not Tbxta is likely to act in endogenous endoderm as discussed later. Tbxta has also previously been shown not to induce ectopic *sox32* when overexpressed with *mixl1* and *gata5* [10].

**Figure 5.**
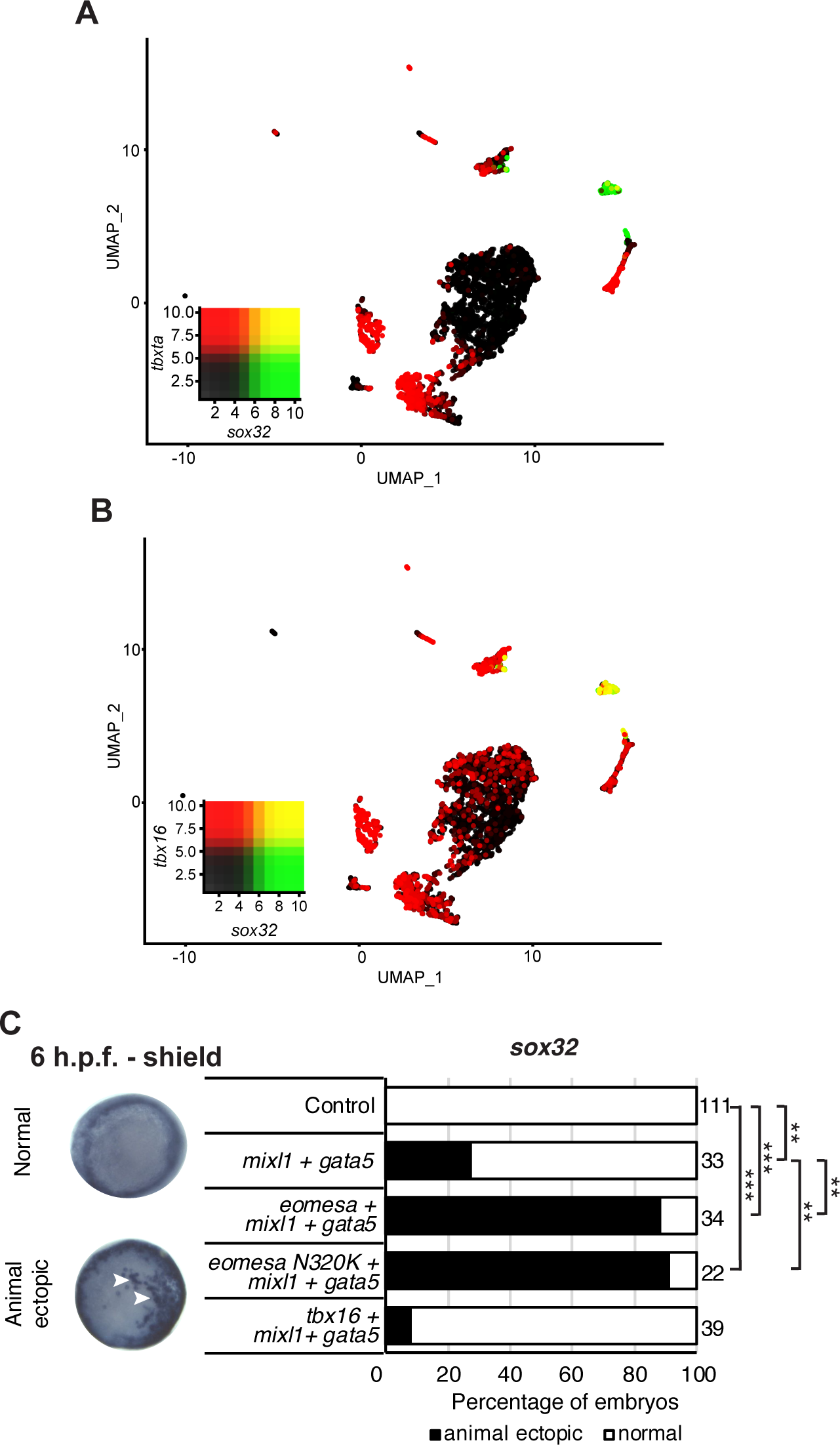
Tbx16 is substantially co-expressed with *sox32* but cannot induce it in combination with *mixl1* and *gata5*. (A-B) UMAP clustering plots of whole embryo single-cell RNA-seq data at shield stage (6 h.p.f.) indicating co-expression of *sox32* with *tbxta* and *tbx16*. Heatmap insets indicate overall expression levels per gene and co-expression. Overlapping expression is shown in yellow. (C) WISH analysis of the ability of *eomesa, eomesa*N320K and *tbx16* in combination with *gata5* and *mixl* to induce *sox32* expression at the animal pole of shield stage (6 h.p.f.) embryos. N = 2. Total numbers of embryos scored per condition are indicated. Representative images of expression patterns per gene per category are shown. Animal views; dorsal to the right. Arrowheads indicate ectopic expression. Fisher’s Exact two-tailed probability test P values: ** *P* ≤ 5×10^−6^; *** *P* ≤ 5×10^−12^.

As expected, combined overexpression of *eomesa, mixl1* and *gata5* induces ectopic *sox32* expression at the animal pole (Fig 5C). The *eomesa* N320K mutation did not markedly influence induction of ectopic *sox32* expression suggesting that this mutation within the T-box does not interfere with the known T-box interactions with Mixl1 and Gata5 [10]. However, *tbx16* did not synergise with *mixl1* and *gata5* to upregulate *sox32* suggesting that Tbx16 and Eomesa are not equally capable of forming a complex with *mixl1* and *gata5* to induce endoderm and/or DFC fate. We conclude that Eomesa and Tbx16 perform similar functions in overlapping processes in the developing zebrafish embryo, but are unlikely to do so through identical molecular mechanisms.

### Eomesa regulates vgll4l expression and DFC formation through interlocking feedforward loops via sox32

Results above demonstrate that Eomesa overexpression induces DFC fate based on both ectopic *sox32* and *vgll4l* expression (Fig 4B,C). To resolve the conflict between the observed Eomesa-mediated repression of *vgll4l* expression at 4 h.p.f. but induction at 6 h.p.f. we explored the role of Eomesa target Sox32 in *vgll4l* induction. We found that *sox32* overexpression through one-cell stage mRNA injection led to a dramatic upregulation of *vgll4l* expression at 6 h.p.f. (Fig 6A), suggesting localised Eomesa-mediated upregulation of *vgll4l* at the margin occurs through Sox32 rather than a switch in Eomesa function directly at the *vgll4l* locus. To test whether Sox32 is required for induction of *vgll4l* expression we performed knockdown using a previously validated and widely used antisense morpholino [38]. This knockdown clearly resulted in loss of DFC *vgll4l* expression (Fig 6B). Furthermore, *vgll4l* induction caused by combinatorial *eomesa*/*mixl1*/*gata5* overexpression was profoundly abrogated by Sox32 knockdown. Thus, during gastrula stages induction of *vgll4l* is via Sox32 rather than direct Eomesa activities.

**Figure 6.**
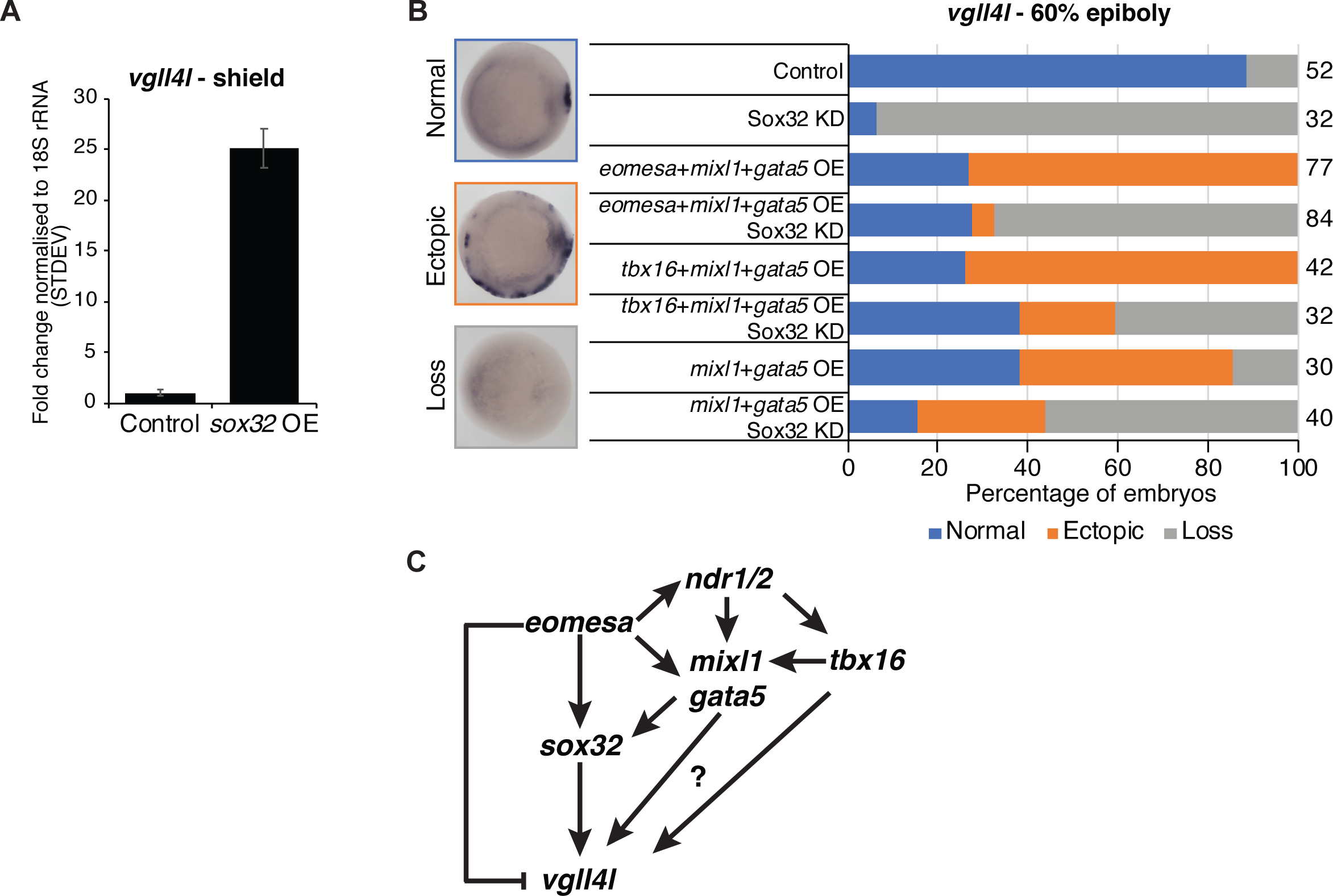
Eomesa activation of *vgll4l* is through feedforward loops via *sox32* and its upstream activators. (A) qRT-PCR analysis of *vgll4l* expression at 6 h.p.f. in control embryos and those injected with *sox32* mRNA at the one cell stage. Expression is represented as fold change relative to control normalised to 18S rRNA. (B) WISH analysis of the ability of *vgll4l* expression at 6 h.p.f. in embryos injected with mRNAs at the one-cell stage as indicated, with and without Sox32 morpholino knockdown. N = 2. Total numbers of embryos scored per condition are indicated. Representative images of expression patterns per gene per category are shown. Animal views; dorsal to the right. (C) Model for Eomesa regulation of *vgll4l* expression in dorsal forerunner cells indicating a type 3 incoherent feedforward loop on the left as Eomesa represses *vgll4l* directly while activating via Sox32, and a potential type 1 coherent feedforward loops on the right as Eomesa activates positive regulators of *sox32* and potentially also *vgll4l*.

*Vgll4l* expression was similarly upregulated by combinatorial *mixl1*/*gata5* overexpression, and partially blocked by Sox32 KD (Fig 6B). However, *mixl1*/*gata5*-induced ectopic *vgll4l* was not as profoundly reduced on Sox32 KD as was the case for *eomesa*/*mixl1*/*gata5* overexpression. It is not completely clear whether this is because *mixl1*/*gata5* can activate *vgll4l* independent of Sox32 function, or due to the absence of Eomesa-mediated repression of *vgll4l*. Addition of *tbx16* to *mixl1*/*gata5* overexpression, had only a small effect on *vgll4l* expression, suggesting *tbx16* does not potently interact with *mixl1*/*gata5* as *eomesa* does nor individually exert a strong influence at the *vgll4l* locus in spite of intron 1 binding (Fig 6B, 4D).

Overall the present results combined with previous published reports from ourselves and others suggest a model wherein Eomesa acts within interlocking incoherent type 3 and coherent type 1 feedforward loops [39] to repress *vgll4l* while combining with Nodal downstream effectors Mixl1 and Gata5 to activate *sox32*, which in turn activates *vgll4l* around the time of DFC specification. In addition to this, our analyses indicate that both mouse Eomes FL and ΔVR isoforms are functionally equivalent to Eomesa, suggesting phenotypic differences between zebrafish and mouse Eomes loss-of-function mutants are not likely to be driven by functional divergence, but rather redundancy with co-expressed factors in zebrafish such as Tbx16.

## Discussion

### Phenotypic differences between mouse and zebrafish Eomes loss-of-function mutants is not due to molecular divergence

T-box transcription factors are an ancient family of genes with many key roles in embryogenesis and disease. Lineage-specific differences that occurred in the family during vertebrate evolution have resulted in altered gene complements and diversity of splice isoforms in distinct evolutionary lineages [1, 40]. While AS events in specific evolutionary lineages have led to functional diversification of certain T-box factors, we have shown that Eomes loss-of-function phenotypic differences between mouse and zebrafish are unlikely to be due to evolutionary differences in Eomes protein function, but rather a degree of compensation by Tbx16 which is present in zebrafish but not mammals.

Though AS is an evolutionary means of increasing functional diversity within the proteome, our data suggests that Eomes exon 6 AS is not functionally important in the context of early development. In the case of the ΔVR splicing event a synonymous mutation created an alternative splice acceptor, however, our data suggests it is hardly used leading to majority production of the FL isoform. That the ΔVR isoform arose and is maintained in the tetrapod lineage may be due to functional equivalence of FL and ΔVR isoforms, leading to a lack of selective pressure.

In overexpression studies the ΔVR isoform has the ability to induce trophoblast markers in embryonic stem cells [41], and cardiac mesoderm markers in embryonal carcinoma cells [7]. Moreover, both FL and ΔVR isoforms can induce organizer markers while repressing ectoderm markers on overexpression in zebrafish. These observations provide further evidence of their functional equivalence.

The present data demonstrate that the ΔVR and ΔCTD isoforms are only weakly expressed compared with FL Eomes. While we find no evidence for the functional importance of the VR it is intriguing that it is both highly conserved and known to be phosphorylated [34]. It is of course possible that these isoforms potentially make substantial contributions in contexts we have not explored. The present evidence, however, suggests that in mice, as in zebrafish the FL isoform is the more important molecule.

### Functional similarities and differences of Eomesa and Tbx16

We previously demonstrated that Eomesa and Tbx16 display overlapping genomic binding profiles in early zebrafish embryos [33]. Whether they are functionally redundant, however, had not been explored. The present experiments strongly suggest that Eomesa and Tbx16 redundantly regulate the homeodomain transcription factor *mixl1*, which has key conserved functions in endoderm formation in zebrafish and mouse [42, 43]. It seems likely that Tbx16 partially compensates for loss of Eomesa during zebrafish endoderm formation. Our study therefore highlights a consistent requirement for T-box function in vertebrate endoderm formation. Interestingly, while multiple orthologous T-box factors have similar expression domains in early zebrafish and mouse embryogenesis, those domains are typically expanded in zebrafish [44]. Coupled with its rapid rate of development and the greater number of T-box factors in zebrafish, there is likely to be a higher degree of T-box factor co-expression, enhancing the probability of redundancy.

While Eomesa and Tbx16 share some redundant functions we also identified key differences. It was previously shown that Eomesa can combine with Mixl1 and Gata5 to drive expression of *sox32* at the animal pole [10]. However, Tbx16 does not appear to have the same ability as Eomesa to drive *sox32* expression either individually or in combination with Mixl1 and Gata5, even though *sox32* expressing cells appear to exhibit *tbx16* expression in single-cell RNA-seq data. This is consistent with previous observations that MZ*eomesa* mutants have reduced expression of *sox32* during gastrulation without complete loss [17]. It therefore seems likely that Tbx16 is sufficient to rescue certain Eomesa functions but cannot completely compensate for its loss. It is further notable that Tbx16 does not seem able to induce the dorsal mesoderm marker *noto* as Eomesa can. However, given that Eomesa appears to act upstream of Nodal [45] it seems likely that major differences in outcome between *eomesa* and *tbx16* overexpression stem from enhanced Nodal signalling on *eomesa* overexpression. It will be interesting to learn more about the common and unique functional activities of Eomes and Tbx16 that drive target gene expression.

Eomesa and Tbx16 are only distantly related within the T-box family (25.3% of Tbx16 amino acid identity), with the majority of conserved amino acids occurring within the T-box domain. Whether they are likely to act in similar protein complexes to regulate their target genes is therefore unclear. A key study in *Xenopus*, however, suggested the specificity of target gene induction is primarily mediated by the T-box itself, rather than NTDs and CTDs [37]. The same study demonstrated a single asparagine to lysine substitution in *Xenopus* Eomes and VegT T-box domains, alter their inductive properties to mimic Brachyury [37]. Importantly, both Eomesa and Tbx16 (which has been proposed as the zebrafish orthologue of *Xenopus VegT* [46]) share the same critical asparagine. Our data suggest that the N320K mutation has little effect on induction of Eomesa target genes explored here, is unlikely to prevent T-box interaction with co-factors Mixl1 and Gata5, or substantially account for differences with Tbxta in endoderm and DFC formation. In fact, analysis of single-cell RNA-seq data suggests that the greater importance of Eomesa and Tbx16 in endoderm formation is more likely to be attributable to Tbxta not being appreciably expressed in the endoderm. It is therefore possible that Eomesa and Tbx16 also have overlapping roles in endoderm formation downstream of driving *mixl1* expression in presumptive endoderm that are yet to be elucidated.

Interestingly, participation of Tbx16 in processes controlled by Eomes in mice is not limited to endoderm formation. For example, while Eomes acts upstream of basic helix-loop-helix transcription factor gene *Mesp1* to specify cardiac mesoderm in mice [7], Tbx16 regulates the orthologous gene *mespaa* in zebrafish [47]. Eomes loss-of-function leads to aberrant mesoderm cell migration during mouse gastrulation, while *tbx16* zebrafish mutants also exhibit cell-autonomous defects in mesoderm migration [48]. It will be interesting to see whether processes known to be controlled by Tbx16 in zebrafish, such as haematopoiesis are regulated by Eomes or other T-box factors in mice [49].

The present study focuses on early embryonic development, however, Eomes is known to have later roles in neurological development, as well as in the immune system. Importantly, Eomes is an key regulator of neurogenesis in the subventricular zone, and loss leads to microcephaly and severe behavioural defects [9]. Though *eomesa* is equivalently expressed in the telencephalon of developing zebrafish larvae, whether null mutants have an equivalent phenotype is unknown [17, 18]. If they do not, however, it is unlikely to be due to redundancy with *tbx16*, which is absent from the developing brain. Similarly, it is unclear whether *eomesa* mutants exhibit defects in the immune system, such as in T cell differentiation and NK cell development and function as in mammals [50, 51]. Both *eomesa* and *eomesb* are co-expressed in lymphocytes in fish, however, suggesting they may be redundant in the immune system [14, 15].

While T-box factor redundancy during development is not a novel concept e.g. [26, 33, 47, 52, 53], the molecular basis for this redundancy (or indeed T-box factor molecular interactions in general) is not well understood. In future it will be interesting to study whether redundant T-box factors recruit similar co-factors to regulate gene expression, and whether this occurs through conserved or divergent amino acid sequences and structural motifs.

### Eomesa, Tbx16, Mixl1 and Gata5 activities during DFC formation

Loss of Eomesa leads to upregulation of *vgll4l* during blastula stages whereas overexpression of *eomesa* causes repression of *vgll4l* [12]. The present experiments suggests that Eomesa acts within feedforward loops to repress *vgll4l* expression until activators including Sox32 accumulate to drive *vgll4l* in DFCs at the onset of gastrulation. Given that Eomesa is maternally contributed and not spatially restricted in early development [17], while accumulation of *vgll4l* activators is principally driven by Nodal at the dorsal margin, this suggests a model wherein Eomesa controls the specificity and timing of *vgll4l* induction. *Eomesa* mRNA steadily declines during blastula stages and is virtually undetectable at the onset of gastrulation, as expression of *mixl1, gata5, tbx16* and *sox32* increase (Fig 3A and [11]). While Eomesa protein does persist through gastrulation [17] it seems likely that temporal and spatial changes in abundance of *vgll4l* activators and repressors acting within these feedforward loops cooperatively regulate the specificity of *vgll4l* expression during DFC specification.

Genetic data, however, suggest that our model is likely to be incomplete. While Sox32 is required for correct DFC formation [54, 55], expression of upstream regulators *mixl1* and *gata5* is not present in DFCs, nor are they required for DFC formation individually or in combination. Rather *mixl1* and *gata5* seem to be strictly required upstream of *sox32* for correct endoderm formation [42, 56, 57]. While we cannot discount the possibility of *mixl1* and *gata5* expression in precursors of DFCs, present evidence suggests that there are either alternative upstream regulators of *sox32* in DFCs vs. endoderm, or additional redundant factors in DFCs rescuing the requirement for *mixl1* and *gata5*. However, given the apparent requirement for Nodal signalling in DFC formation [58], it seems likely that whatever the upstream regulators of *sox32* expression in DFCs they will be Nodal-dependent. Overall this highlights a lack of understanding of the gene regulatory networks that direct DFC vs. endoderm formation, which will be a key focus of our future work.

Recent evidence suggests Vgll4l is required for *tbx16* expression during DFC formation [13]. That Tbx16 binds the *vgll4l* promoter during gastrulation could suggest that complex regulatory loops control DFC formation and maintenance. The ability of Eomesa to induce ectopic DFCs during early gastrulation combined with expression of mouse Eomes in progenitors of the node and requirement for correct node formation [6, 7] suggests the potential for a conserved role in establishment of left/right asymmetry with some degree of redundancy with Tbx16 in zebrafish. However, a role for the *vgll4l* mammalian homologue *Vgll4* in left/right asymmetry has yet to be determined. Further study of the mechanistic parallels in T-box factor mediated formation of zebrafish DFCs and mouse node would be beneficial to gain a more detailed evo-devo understanding of this process.

Overall we conclude that enhanced AS in mammals has not significantly altered Eomes function in early embryogenesis. Rather we conclude that the different degrees of T-box factor co-expression and the presence/absence of additional factors including Tbx16 has modulated the severity of the Eomes null mutant phenotype in the embryo proper between mouse and zebrafish. Furthermore, we conclude that in zebrafish Eomesa participates in DFC formation through directing feedforward loops via *sox32* to control *vgll4l* expression. Our results therefore provide novel insights into evolutionary differences in vertebrate endoderm formation, and the gene regulatory networks involved in controlling the zebrafish left-right organiser formation.

## Materials and Methods

### Zebrafish strains

AB and mutant zebrafish were reared as described [59]. For *eomesa* mutant experiments eggs from homozygous *eomesa*^*fh105/fh105*^ females were *in vitro* fertilized with *eomesa*^*+/fh105*^ sperm yielding a mixture of M*eomesa* and MZ*eomesa* mutant embryos. Since previous studies have revealed no differences in endodermal or mesodermal expression between M*eomesa* and MZ*eomesa* mutant embryos we did not distinguish between them in this study [17, 45]. All zebrafish studies complied fully with the UK Animals (Scientific Procedures) Act 1986 as implemented by King’s College London, The University of Warwick, or were in accordance with the policies of the University of Toronto Animal Care Committee.

### Cloning for in vitro production of mRNAs and mammalian expression vectors

Full length *tbx16* and *eomesb* open reading frames were cloned with C-terminal myc tags into XhoI and XbaI sites in pCS2+ by PCR from zebrafish cDNA using the following primers: *tbx16-myc* CATACTCGAGATGCAGGCTATCAGAGACCT and CGCGTCTAGACTACAGATCCTCTTCTGAGATGAGTTTTTGTTCCCAGCACGAGTATGAGAAAA; *eomesb-myc* ATATCTCGAGATGCCCGGAGAAGGATCCAG and GCGCTCTAGACTACAGATCCTCTTCTGAGATGAGTTTTTGTTCGCTGCTGGTGTAGAAGGCGTA. Full length *gata5* cDNA with a C-terminal HA tag was similarly cloned into pCS2+ EcoRI and XhoI sites using primers CGCCGAATTCATGTATTCGAGCCTGGCTTT and AATGCTCGAGTCAAGCGTAATCTGGAACATCGTATGGGTACGCTTGAGACAGAGCACACC. Eomes cloning into pCS2+ was performed between EcoRI and XhoI sites. pCS2+*eomesaN320K* was produced by PCR mutagenesis of the wild type construct using AAACTGAAGCTAACCAACAAGAAAGGAGCAAATAACAACAAT and TCCGAAAGATATTTCTTGTCT followed by recircularization. *Eomesa* ΔCTD was similarly produced using the following primers TAAGAACTGCTTTTCAAGATCCTTTATCAATCC and CGAATCATAATTGTCCCTGAA. The ΔNTD mutation was produced by removing the EcoRI/BstEII fragment from pCS2+*eomesa* and replacing with the EcoRI/BstEII fragment produced by PCR from from pCS2+*eomesa* with primer pair GCCCTCGAATTCACAGTTAAGAATGGCGCGGGCGC and CCCGCAGGTCACCCACTTTCCGCCCTGAAATCTCCA.

### mRNA, morpholinos and microinjections

All capped mRNA were synthesized from plasmids encoding proteins of interest in pCS2+ NotI linearization followed by SP6 transcription as described [11], with the exception *tbxta* which was produced from pSP64T as described [60]. mRNA quantities for T-box factors were scaled in order to inject equimolar amounts of each mRNA per embryo. One-cell stage embryos were injected with the following quantities: *eomesa* – 400pg; *EomesΔVR* – 410pg; *EomesFL* – 420pg; *eomesaΔNTD* – 308pg; *eomesaΔCTD* – 285pg; *eomesaN320K* – 400pg; *eomesb-myc* – 286pg, *tbx16-myc* – 217pg; *tbxta* – 223pg; *gata5-HA* – 140pg; *mixl1* – 200pg. For Tbx16 knockdown one-cell stage embryos were injected with, 0.5 pmol of a previously characterized *tbx16* morpholino (GeneTools)[61].

### In vitro protein production

Unlabelled *in vitro* translated protein was produced using rabbit reticulocyte lysates according to manufacturer’s protocol (Promega).

### Northern blot

Total RNA was extracted from specified cell types using Rneasy Mini Kits (QIAGEN), and polyA selected using Oligotex mRNA Mini Kits (QIAGEN). 500ng polyA+ RNA per lane was size fractionated on a 1.5% agarose/MOPS gel, transferred onto Hybond N membranes (GE Healthcare), and probed with ^32^P-random-primed 1kb XmaI-EcoRV cDNA fragment corresponding to the exon 1-4 T-box region.

### Western blot

Cell lysates were prepared using radioimmunoprecipitation assay (RIPA) buffer, subjected to SDS– polyacrylamide gel electrophoresis and transferred onto polyvinylidene difluoride membranes. Membranes were blocked with 5% milk powder in Tris-buffered saline with Tween 20, incubated in primary antibodies overnight including rabbit anti-Eomes CTD (Abcam, ab23345, 1:2,000), rabbit anti-Eomes NTD (Santa Cruz, sc-98555, 1:1000) and rat anti-Eomes (eBioscience, 14-4876, 1:1000). Secondary antibodies were donkey anti-rabbit horseradish peroxidase (GE Healthcare NA934, 1:2,000) and goat anti-rat horseradish peroxidase (GE Healthcare NA935, 1:2,000). Blots were developed by chemiluminescence using Amersham ECL Prime Detection Reagent (GE Healthcare).

### Embryonic stem cell differentiation

Wild type (+), *Eomes*^*null/null*^ (null), feeder-depleted ESCs were cultured in DMEM (ThermoFisher) with 15% FCS, 1% non-essential amino acids, 0.1 mM β-mercaptoethanol and 1,000 U/ml recombinant leukaemia inhibitory factor (Millipore). For differentiation ESCs were resuspended at 1×10^4^ cells/ml in DMEM (ThermoFisher) with 15% FCS, 1% non-essential amino acids, 0.1 mM β-mercaptoethanol in hanging drops (10 μl) plated on the inside lids of bacteriological dishes. After 48 h embryoid bodies were transferred in 10 ml medium to 10 cm bacteriological dishes and RNA extracted at the appropriate timepoints.

### P19Cl6 cell culture and differentiation

P19Cl6 embryonal carcinoma cells were cultured in α-MEM (ThermoFisher) supplemented with 10% FCS. To induce differentiation, cells were seeded at 3.7×10^5^ cells/6 cm dish in media containing 1% DMSO (Sigma) and RNA harvested after 72 hours.

### Cytotoxic T-cell lymphocyte (CTLL), SNH fibroblast and HeLa cell culture

CTLL cells derived from the ATCC TIB-214 line were maintained at 10^4^-10^5^ cells per ml in complete T cell medium supplemented with IL-2. SNH fibroblasts and HeLa cells were maintained on 0.1% gelatin coated dishes in DMEM supplemented with 10% bovine calf serum (Hyclone).

### RT-PCR

Cytoplasmic RNA was produced as previously described [62]. Total RNA was produced using Rnasy Mini Kits according to manufacturers protocol (QIAGEN). RT-PCR was performed using OneStep RT-PCR Kit (QIAGEN) using the following primers: Total Eomes – TGTTTTCGTGGAAGTGGTTCTGGC and AGGTCTGAGTCTTGGAAGGTTCATTC; Eomes exon 4-6 to distinguish FL and ΔVR isoforms ATCGTGGAAGTGACAGAGGACG and CGGGAAGAAGTTTTGAACGCC; Gapdh – TGCACCACCAACTGCTTAGC and GGCATGGACTGTGGTCATGAG; Eomes start codon to ΔCTD 3’ UTR – ATATCTCGAGATGCAGTTGGGAGAGCAGCTC and TGGGCTCGAAGATGAAACTC; *HBB* exon 2 to Eomes exon 6 – GCACGTGGATCCTGAGAACT and CGGGAAGAAGTTTTGAACGCC. For nested PCR to test exon 5-6 splicing association with the long Eomes 3’ UTR the initial primer pair used was ATCGTGGAAGTGACAGAGGACG and CAAGTACGGAGGCAGCTGAG.

### Whole-mount embryo staining

Whole-mount *in situ* hybridization (WISH) of zebrafish embryos were performed as described [63]. Anti-sense riboprobes for *noto* [64], *chd* [65], *vgll4l* [12], *zic3* [66], *mixl1* [55] and *sox32* [38] were produced as described.

### Cloning and mutagenesis to test Eomes exon 6 splice sequences

Clones to test the splicing efficiency at *Eomes* exon 6 were generated by cloning PCR products using primers ACGGCAATTGGCCTCGAACATTCTTGCTTC and CCAGCCATCACTTTGGTCAAAGGTGGAAGGCAAAAG into MfeI-BstXI sites of the human *HBB* gene (GenBank accession no. U01317) within a previously described TAT-inducible expression vector [67]. Mutation of splicing sequences for the ΔVR and FL Eomes isoforms were introduced by PCR using primers: ΔVR - CATGTACACGGCTTCAGAAAACGACAGGTTAACGCCAAGTCCGACGGATTCCCCTCGATCCCATCAGATTGTCCCTGG and CTACAATATAAAGAGAGACACTTAAAAATAAAAAACAACCCTCACGTTGTCCCCAAACAAGCTGCCTCCCAGAAGC; FL – CATGTACACGGCTTCAGAAAATGACAGGTTAACTCCATCTCCCACGGATTCCCC and GGACATTATATACACCGCCTCTTATATTTTTACACCAACCCTCACGTTGTCCCCAAACAAGCTGCCTCCCAGAAGC followed by recircularization of the resulting PCR products. Deletion of the VR was similarly performed using primers ATCCCATCAGATTGTCCCTGGA and CTACAATATAAAGAGAGACACTTAAAAATAAAAAACAACCCTCACGTTGTCCCCAAACAAGCTGCCTCCCAGAAGC. HBB plasmids were co-transfected with a plasmid expressing the HIV transactivator protein TAT [68], into HeLa cells using Lipotectamine 2000 according to manufacturers protocol (ThermoFisher).

### Conservation analysis

The Tbr1 subfamily Gene Tree was generated by Ensembl [69]. *Eomes* conservation measurements (*phyloP*) across 60 vertebrate species were visualized in UCSC Genome Browser (http://genome.ucsc.edu/) [70, 71]. Sequence logos of the Eomes variant region in placental mammals, other tetrapods and teleosts were based on alignment of the same 60 vertebrate species and visualized using WebLogo [72].

Full-length protein alignments were performed using Clustal Omega [73-75] and visualized using JalView [76]. BLOSUM62 average distance gene tree was also produced using Jalview.

### Single-cell RNA sequencing analysis

Single cell mRNA sequencing count data of zebrafish embryonic cells from Wagner *et al* (2018) was downloaded from GEO [36, 77]. Raw UMI-filtered count data in CSV form from the untreated embryos at 6 hours post fertilisation (GSM3067190) was imported in to R v3.6.3 and analysed using Seurat v3.1.2 [78]. Cells with less than 200 features, and features detected in <3 cells were discarded. The remaining count data were then normalised via SCTransform v0.2.1 [79] with mitochondrial genes passed as a regression variable. Genes were clustered using UMAP utilising the R package uwot v0.1.5 [80], with the following parameters: dims = 1:30, n.neighbors = 5, min.dist = 0.001.

## Authors’ contributions

Conceived and designed the experiments: ACN. Performed the experiments: ACN, CDT, AEEB, SJC, FCW and RE. Analysed the data: ACN, CDT, MDW. Wrote the paper: ACN.

## Acknowledgments

We thank Elizabeth Bikoff and Elizabeth Robertson for reagents, valuable conversations and critical review of the manuscript. We also thank Eirini Vlachaki for technical assistance, Claire Simon, Ita Costello, Arne Mould, Mick Dye, Nick Proudfoot, Kacper Wozniak, Karuna Sampath and Daniel Hebenstreit for valuable discussions, the Warwick zebrafish facility for zebrafish care and acknowledge the contributions of Henrietta Lacks and her family to this research.

