## Supplementary Figures 1-2 for "Eomesodermin is functionally conserved between zebrafish and mouse in spite of different mutant phenotypic severities, and controls left/right organiser formation via interlocking feedforward loops"

### Figure S1

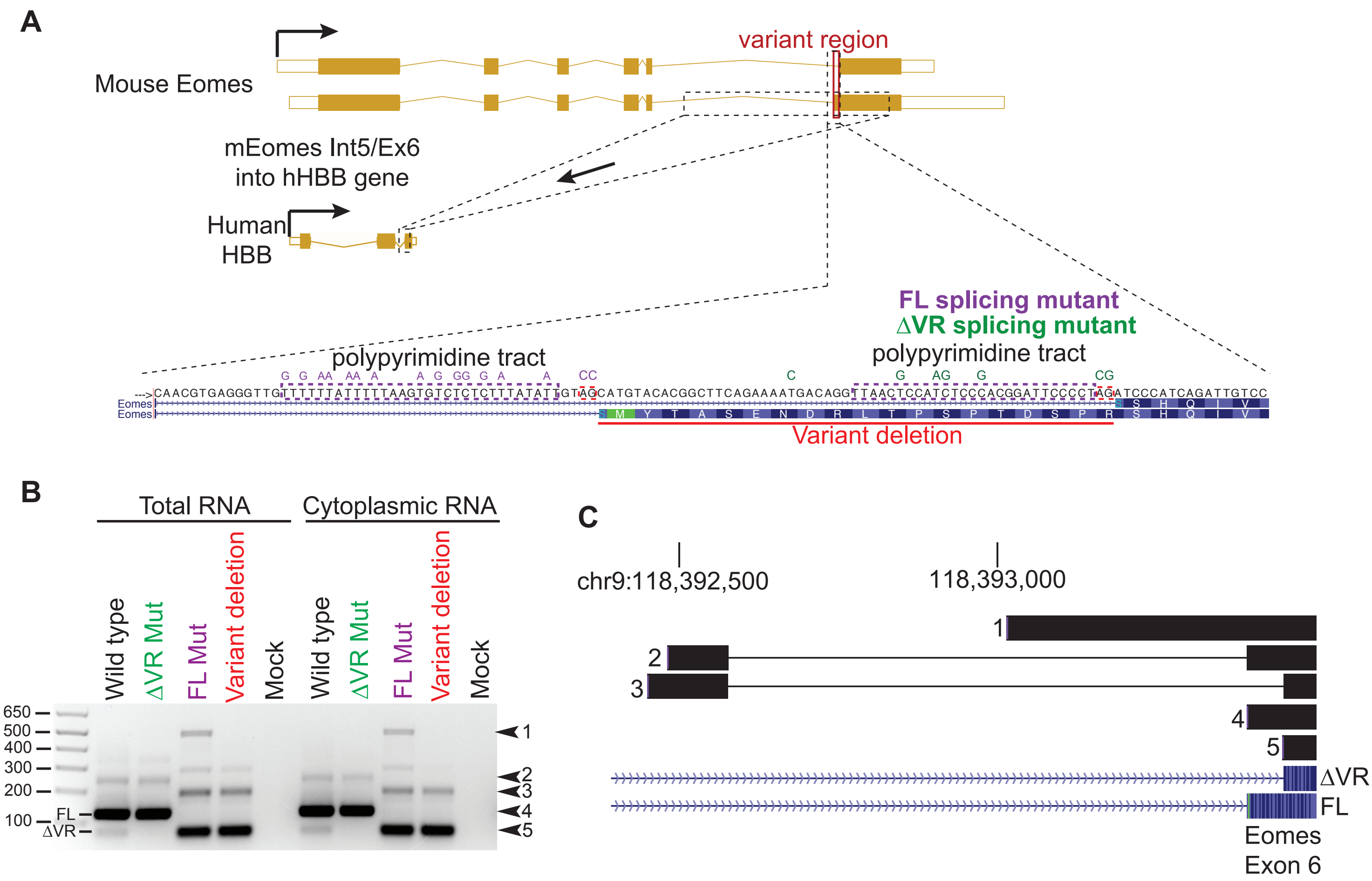


**Figure S1. Mutational analysis of exon 6 splicing sequences indicate that the *Eomes* ΔVR splice site is weak compared to the *Eomes* FL splice site.**

(A) Schematic of hybrid gene construction and mutagenesis to investigate FL and ΔVR exon 6 splicing consensus sequences. *Eomes* intron5/exon 6 sequences cloning substitution into the body of *HBB* is indicated. Base mutations or deletions in each construct are colour coded. All mutations within the VR are synonymous.

(B) RT-PCR revealing HBB exon 2/Eomes exon 6 splicing events in HBB/Eomes hybrid genes. Arrows indicate sequenced PCR products. Note that the ΔVR splicing event is uncommon, reflecting other results. Further note that mutation of ΔVR splicing sequences leads to all splicing into the FL site, while mutation of the FL splicing sequences leads to discovery of intron 5 cassette exons rather than complete splicing into ΔVR, potentially reflecting weakness of ΔVR splicing sequences.

(C) Genomic alignments of RT-PCR products labelled in panel B.

#### Figure S2

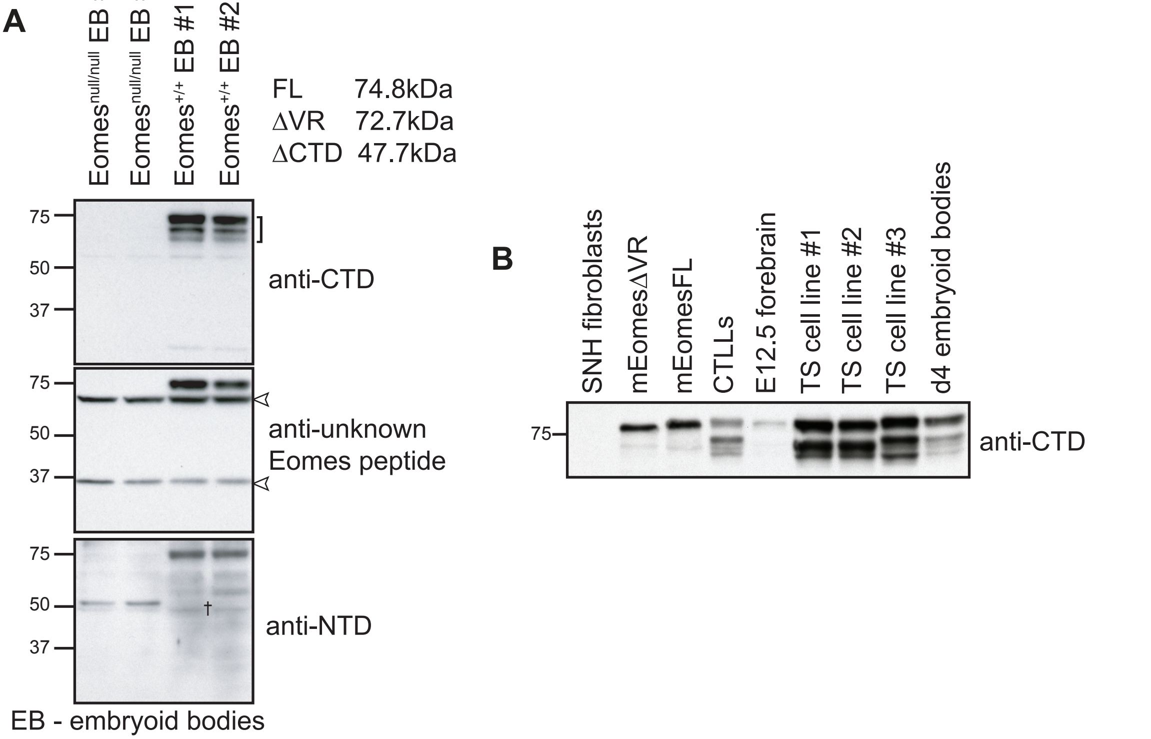


**Figure S2. Analysis of *Eomes* protein products indicates lack of ΔCTD isoform and multiple N-terminal deletions.**

(A) Western blot analysis of Eomes null and wild type day 4 embryoid bodies using antibodies recognizing different regions of the Eomes protein. Bracket indicates Eomes bands identified by anti-CTD antibody. Open arrowheads indicate non-specific bands. † indicates potential ΔCTD isoform band.

(B) Western blot analysis of different Eomes expressing cell types. SNH fibroblasts are to control for background signal.
